# *PEAPOD* limits developmental plasticity in *Arabidopsis*

**DOI:** 10.1101/102707

**Authors:** Derek W. R. White

## Abstract

Higher plants utilise developmental plasticity to adapt to changes in the environment, especially to variations in light. Much of this change in growth and development involves the light-mediated regulation of multiple hormone pathways. However, despite considerable progress towards understanding the molecular processes controlling light signalling and hormone activity, regulatory mechanisms preventing exaggerated plant developmental responses are not well understood. Here I report that the *PPD* regulatory complex has a crucial role in limiting developmental plasticity in *Arabidopsis*. Reductions in *PPD* or *KIX8/9* gene expression resulted in; tolerance to ABA inhibition of seed germination, hypocotyl elongation, increases in stomata on hypocotyls, cambial cell proliferation and seed weight, and delayed flowering. Transcript profiling and analyses of hormone responses and genetic interactions established PPD modulates developmental plasticity, mainly by a combination of transcriptional activation and repression of genes controlling CRY/PHY light signalling and ABA, auxin, brassinosteroid, cytokinin and gibberellin homeostasis.

## Introduction

Because they are sessile, higher plants have evolved extensive developmental plasticity to acclimatize to changes in the environment. This capacity to alter plant morphology involves coordinated changes in cell proliferation and expansion during growth and development, throughout the life cycle. Light is one of the main environmental signals influencing plant developmental plasticity. In addition to its importance for photosynthesis, light also affects many aspects of plant development including seed germination, seedling de-etiolation, photomorphogenesis, stem elongation, leaf size, stomatal density, shade avoidance, flowering time, and seed development^1–3^. The perception of light by photoreceptors can cause changes in the transcription of ∼30% of genes in the *Arabidopsis* genome, including transcription factors that modulate hormonal, metabolic, stress, pigment biosynthesis, and defence pathways^4–5^.

Plants perceive light using multiple photoreceptors: in *Arabidopsis* there are five red/far red absorbing phytochromes (phyA – phyE), two blue absorbing cryptochromes (cry1 and cry2), two blue/UV absorbing phototropins (phot1 and phot2), and a UV-B receptor, UVR8^6–7^. Although individual photoreceptors elicit specific developmental responses, light signals mediated by different photoreceptors are also integrated and can have similar effects on transcription and morphology during seedling photomorphogenesis. The integration point for multiple light receptors appears to be a few light-induced transcription factors first identified as positive regulators of seedling photomorphogenesis, one of which is the bZIP transcription factor, HY5^8–11^. HY5 acts downstream of the phytochrome, cryptochrome, and UV-B photoreceptors, directly regulating expression of ∼3,800 genes in the *Arabidopsis* genome including genes involved in photosynthesis, light signalling, hormone biosynthesis and catabolism, circadian rhythm, flavonoid accumulation, and nutrient assimilation^9^. In the dark or shade a negative regulator of photomorphogenesis, COP1, a RING-finger type ubiquitin E3 ligase, physically interacts with HY5 and mediates its degradation^12^. HY5 degradation is enhanced by the physical interaction of COP1 with SPA proteins^12–13^. When seedlings are exposed to light, activated cry1, cry2, phyA, or phyB directly interact with SPA proteins resulting in, dissociation of the COP1-SPA complex, accumulation of HY5, and promotion of photomorphogenic development^14–17^. This interaction of light, photoreceptors and both negative and positive regulators of light signalling provides a mechanism to fine tune plant growth and developmental responses to changes in light quantity and quality.

Coordinated changes in the homeostasis and signalling of multiple hormones are an essential component of light-mediated alterations in growth and development^3,18^. Hormones such as auxin, brassinosteroid (BR), and gibberellin (GA) antagonise photomorphogenesis, whereas it is promoted by abscisic acid (ABA). For example, light influences auxin levels by coordinated regulation of the transcription of *TAA1*, an auxin biosynthesis gene, and *SUR2*, a cytochrome P450 monooxygenase that alters auxin levels by converting indole acetic acid to indole glucosinolates^18–21^. Active phyB reduces auxin levels by the up-regulation of *SUR2*, and down-regulation of *TAA1* transcription, whereas inactivation of phyB results in elevated levels of auxin due to reduced *SUR2* and increased *TAA1* transcription. Similarly, the light-mediated regulation of BR, GA, and ABA levels are modulated by a combination of metabolic processes; biosynthesis, conjugation, and catabolism^22–34^. However, despite extensive progress made in recent years towards understanding the mechanisms controlling light signalling and hormone activity, one of the fundamental questions in plant biology remains how plants coordinate and optimise growth and development in a constantly changing environment.

Higher plants are able to adapt to changes in the environment by altering the size of their leaves. In *Arabidopsis*, as the leaf grows cells differentiate and undergo cell expansion, starting at the tip of the leaf and progressing to the base^35^. At the same time meristemoids, stomatal lineage precursors, undergo a limited number of asymmetric cell divisions before forming stomatal guard cells and associated pavement cells^36^. Meristemoid cell proliferation is an important determinant of leaf size in *Arabidopsis*, because meristemoid cells generate a large portion of the epidermal pavement cells (48% in leaves and 67% in cotyledons)^37^. To date the only mechanism reported specifically limiting meristemoid cell proliferation involves PPD1 and PPD2, and interacting protein partners^38–39^. Deletion or silencing of the tandemly repeated *PPD* genes in *Arabidopsis* extends division of meristemoids in the epidermis, increasing stomatal density and leaf size, whereas overexpression of *PPD1* results in the opposite phenotype. The PPD proteins belong to class II of the plant-specific TIFY family of transcription regulators. This group of proteins, which includes the JAZ proteins, negative regulators of jasmonic acid signalling, have a central ZIM domain that mediates interactions between TIFY proteins and also with the EAR domain containing NINJA, which acts as an adaptor for the corepressor TPL^40–41^. The PPD proteins are distinguished from other TIFY proteins by a unique N-terminal PPD domain. This domain mediates the interaction of PPD proteins with the EAR domain containing TPL adaptor proteins KIX8 and KIX9^39^. Plants with mutant *kix8* and *kix9* have a similar leaf phenotype to the *Δppd* deletion mutant. PPD proteins also physically interact with SAP, an F-box protein that forms part of a SKP1/Cullin/F-box E3 ubiquitin ligase complex which targets the PPD proteins for degradation^42^. Plants overexpressing *SAP* also have a leaf phenotype similar to the *Δppd* mutant. Hence the PPD proteins and their interacting partners provide a molecular mechanism to change leaf size.

To assess if the *PPD* genes have a more extensive role in the control of plant developmental plasticity I first characterised the phenotypes of *Arabidopsis Δppd* deletion mutant and *PPD1* overexpression genotypes throughout the life cycle. I then used genome-wide transcript profiling to identify genes with altered transcription levels due to *PPD* deletion. Hormone and hormone biosynthesis inhibitor treatments, and genetic analysis were used to confirm alterations in hormone homeostasis detected by transcript profiling. Finally, I used a combination of genetic and gene expression analyses to provide insights into how PPD proteins regulate both hormone homeostasis and light signalling. Here I report an extensive role for the *PPD* genes as master regulators modulating the plasticity of many aspects of development.

## Results

**The PPD regulatory complex controls diverse aspects of growth and development**. To further understand the role that *PPD* genes have in the control of plant development a comparison was made of cell proliferation and growth characteristics in *Arabidopsis* wild-type, *ppd* deletion mutant (*Δppd*), and *Δppd::PPD1* overexpressor (*PPDOE*) genotypes. As previously reported *Δppd* mutant plants had enlarged dome shaped leaves with elongated petioles and larger cotyledons than wild type, whereas overexpression of *PPD1* resulted in plants with smaller leaves and cotyledons, and shorter petioles^38^ (Fig 1a). Seedlings of the *Δppd* mutant had increases in hypocotyl length and stomata numbers on the hypocotyl (Fig. 1a,b,e,f). Surprisingly, *PPDOE* seedlings also had an increase in hypocotyl length. However, stomata numbers on the hypocotyls of *PPDOE* were less than wild type. Additionally, *PPDOE* primary roots were shorter than wild type (Supplementary Fig. 1).

**Figure 1.**
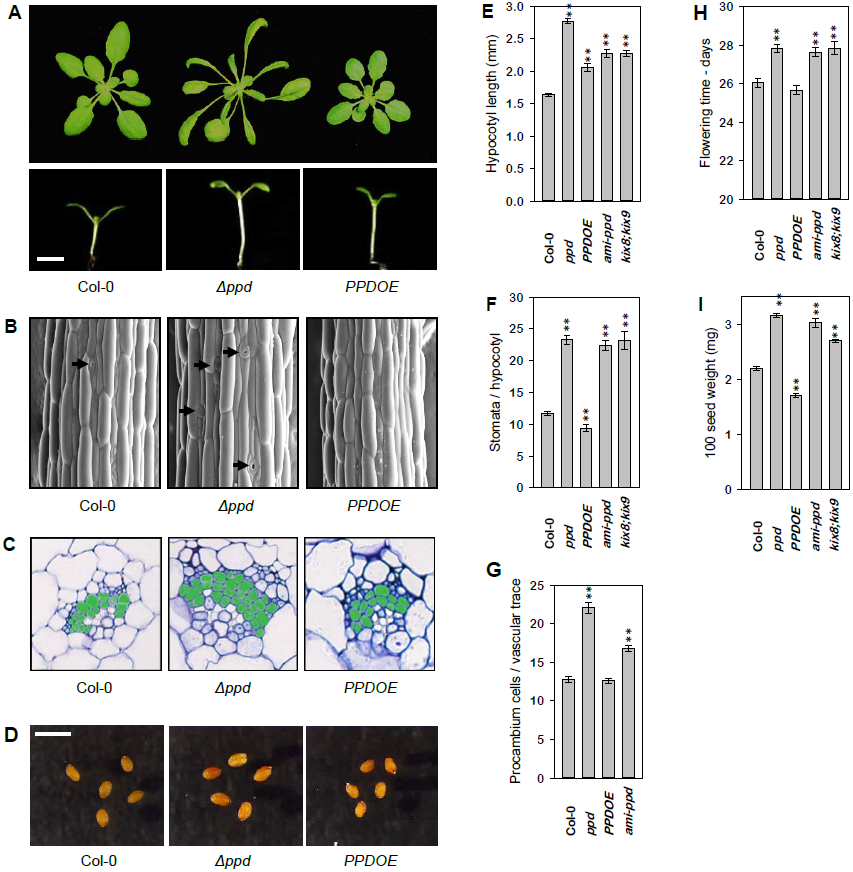
***PPD* has a multifaceted role in plant growth and development.** Comparison of the phenotypes of wild type Col-0, *Δppd* deletion, and *PPDOE* over expression genotypes, (A) mature plants at 21 days post germination (dpg) (upper) and seedlings 5 dpg (lower, bar 1mm), (B) SEM of hypocotyls 3 dpg (bar 100µm), black arrows indicate stomata, (C) transverse sections of cotyledon petiole vascular traces at 10 dpg, with procambium cells shaded green, and (D) representative examples of seed size (bar 1mm). Graphs compare wild type, *Δppd, PPDOE, ami-ppd*, and *kix8;kix9* genotypes for (E) hypocotyl length 5 dpg, (n=30), (F) stomata number per hypocotyl 5 dpg, (n=40), (G) 100 seed weight, (n=7) and (H) days to flowering for plants grown in a 14 hr light:10 hr dark photoperiod, (n=20). (I) Comparison of procambium cell proliferation in the cotyledon petiole vascular traces of wild type, *Δppd, PPDOE* and *ami-ppd* seedlings at 10 dpg, (n=30). \*\**P*<0.001 compared with the wild type (2-tailed Student’s *t*-test). Error bars represent ±SE.

I previously reported that *Δppd* has a more extensive, and *PPDOE* a reduced, vascular network in cotyledons^38^. Histochemical staining of the roots of *PPD1promoter-GUS* plants indicated *PPD1* is expressed within the vascular stele (Supplementary Fig. 2). Furthermore, up-regulation of *CYCB1;1-GUS*, a marker of cell-cycle progression, in both the root and leaf vascular of *Δppd*, suggested a role for *PPD* genes controlling (pro)cambium cell proliferation (Supplementary Fig. 3). This function was confirmed with the observation that *Δppd* had an increase, and *PPDOE* a reduction, in procambium cell number in cotyledon vascular traces (Fig. 1c,g). Similarly, cambium cell proliferation increased in *Δppd*, and decreased in *PPDOE*, mature inflorescence stems (Supplementary Figs. 4 and 5). Furthermore, in a comparison of vascular development in seedling hypocotyls *Δppd* had increased xylem vessel number, whereas in *PPDOE* the amount of xylem was reduced (Supplementary Fig. 5).

I also observed that *Δppd* plants grown in a long-day photoperiod were delayed in flowering (Fig. 1h), whereas *PPDOE* flowered at the same time as wild type. In addition, seed of *Δppd* appeared to be larger than wild type, while seed of the *PPDOE* line were smaller (Fig. 1d). These seed size phenotypes were reflected in significant differences in seed weight, with a 53% increase for *Δppd* and a 22% decrease for *PPDOE* (Fig. 1i).

The role of the *PPD* genes in this multifaceted *Δppd* mutant phenotype was verified in two ways. Firstly, a *PPD1* transgene was sufficient to complement the *Δppd* deletion phenotype (Figs.S5-6). For all the characters examined; hypocotyl length, stomata number per hypocotyl, cotyledon procambium cell number, stem cambium cell proliferation, hypocotyl xylem vessel number, flowering time and seed weight, the *Δppd::PPD1* line was not significantly different from wild type. Secondly, other genotypes with disruption of the *PPD* complex had phenotypes similar to the *Δppd* mutant. A transgenic line with partial silencing of *PPD1* and *PPD2* expression, *ami-ppd*, due to overexpression of an artificial microRNA^39^, had increased stomata numbers on the hypocotyl, delayed flowering time, and an increase in seed weight, similar to *Δppd* (Fig. 1f,h,i), and increases in hypocotyl length and procambium cell proliferation significantly greater than wild type (Fig. 1e,i). Also, in a pattern similar to *Δppd* and *ami-ppd*, *kix8;kix9* mutants had significant increases in hypocotyl length, stomata numbers on the hypocotyl, flowering time, and seed weight (Fig. 1e,f,h,i). The complex phenotype resulting from altered *PPD* or *KIX* gene expression suggests the PPD transcriptional regulators have a role coordinating and controlling diverse aspects of growth and development, throughout the *Arabidopsis* life cycle.

**Genome-wide transcriptomics reveals PPD regulates light signalling and hormone metabolism**. To obtain insight into transcriptional changes underlying the *ppd* deletion phenotype, RNA was isolated from whole seedlings of *Δppd* and wild type plants at 10 days post germination and compared by RNA-Seq genome-wide transcription analysis. A total of 2,830 genes were differentially expressed in *Δppd* compared to wild type, with 927 (33%) up-regulated and 1,903 (67%) down-regulated (Supplementary Table 1). Gene Ontology terms significantly over represented included those for; “response to light stimulus”, “circadian rhythm”, “hormone biosynthesis process”, “regulation of hormone levels”, and “stomatal complex development” (Fig. 2a). To determine which of these genes might be direct targets of the PPD proteins, the dataset was compared with published information on the genome binding sites of PPD2^39^ (Supplementary Table 1). Significant enrichment of PPD2 targets was found among genes both up- and down regulated in the *Δppd* mutant (8.9% or 251, with 30% up-regulated) (Fig. 2d). By chance, PPD2 would be predicted to bind to 3.5% of genes in the dataset (1,191 targets out of 33,602 *Arabidopsis* genes). Consistent with the increased number of cells involved in stomatal development in the *Δppd* mutant, most of the genes known to be involved in regulation of meristemoid cell proliferation were present in the dataset, all up-regulated (Table IA). The initiation and proliferation of meristemoids is regulated by a basic helix-loop-helix transcription factor, SPCH^43^, which has a large number of binding sites in the genome (8,327)^44^. These SPCH targets were cross-referenced with the *Δppd* differential gene expression dataset to determine the extent of targeting by SPCH (Supplementary Table 1). Significant enrichment of SPCH targets (1,050) was found in the dataset (37%), but with fewer up-regulated (12.5%) than expected (Fig. 2d). By chance, SPCH would be expected to bind to 25% of genes in the dataset. There were fewer joint PPD + SPCH target genes in the dataset (26) than expected indicating limited functional overlap between the gene expression networks regulated by these two transcription factors. RT-qPCR expression analysis confirmed the up-regulation of *SPCH, TMM*, and *ERL1* in *Δppd*, while all three genes were down-regulated in *PPDOE* (Fig. 2b).

**Figure 2.**
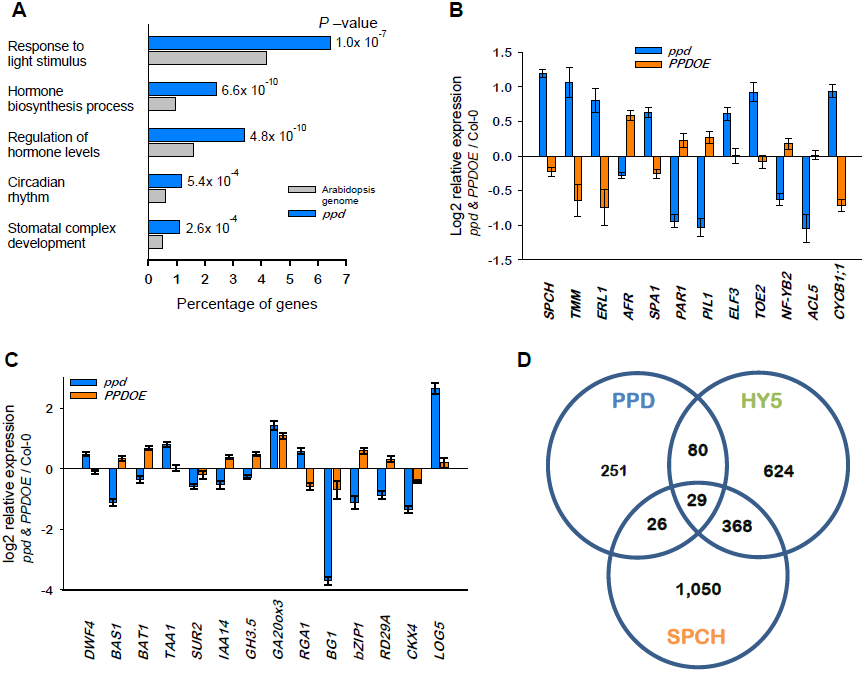
**Genome-wide differential gene expression between wild type and *Δppd* mutant reveals *PPD* has roles regulating light signalling, circadian rhythm, hormone metabolism and stomatal development.** (A) A selected subset of the Gene Ontology terms significantly enriched among the genes identified by RNA-Seq analysis of seedlings as differential expressed in *Δppd*. RT-qPCR expression analysis of selected genes regulating (B) stomatal development, light signalling, circadian rhythm, photoperiodic flowering, and cambium proliferation, and (C) hormone biosynthesis, catabolism, conjugation and signalling genes, compared the *Δppd* mutant and *PPDOE* over expression genotypes relative to wild type. (D) Diagram of the number of genes differentially expressed in the *Δppd* mutant that are putative binding targets of PPD2, SPCH or HY5. Error bars represent ±SE.

**Table 1.**
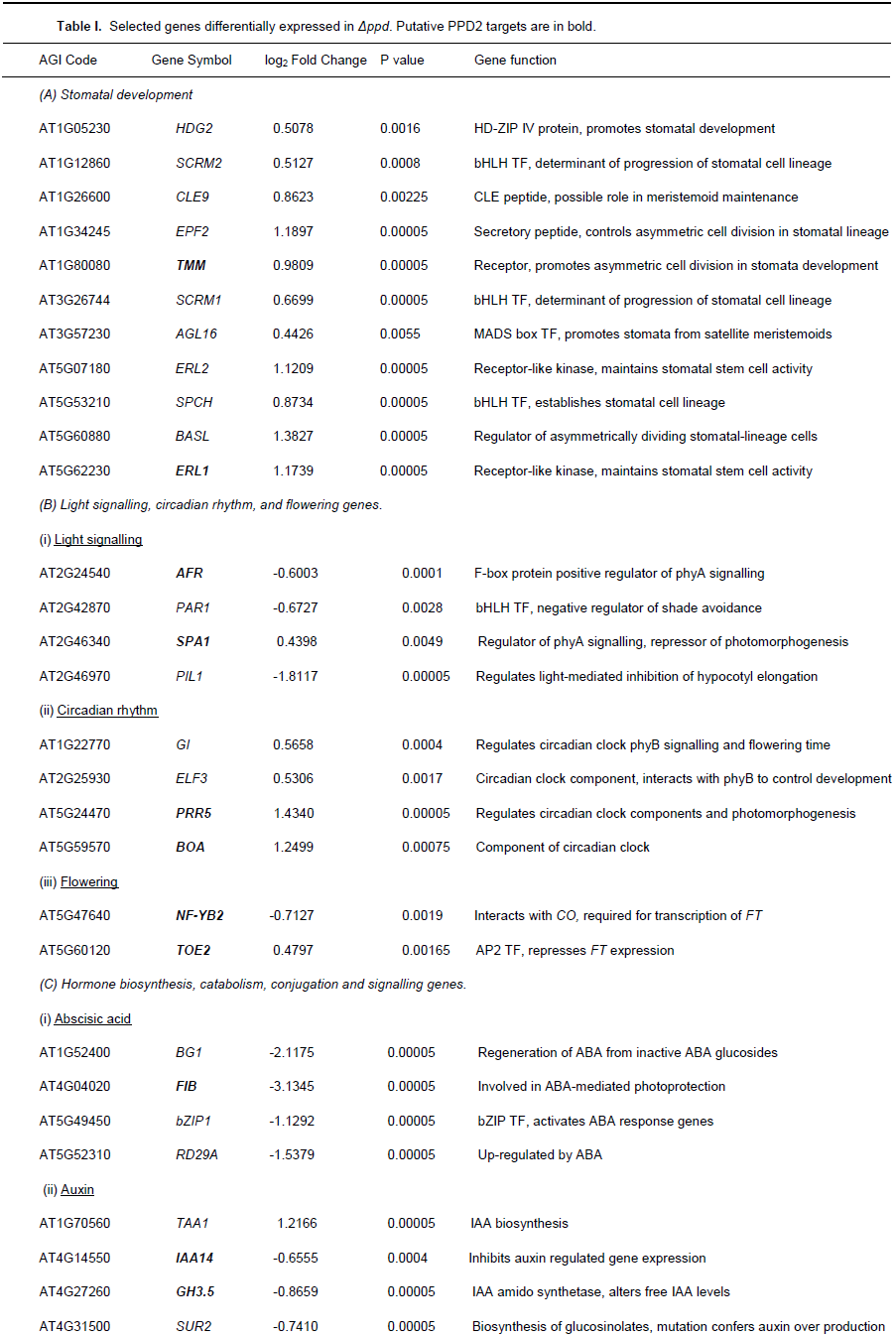
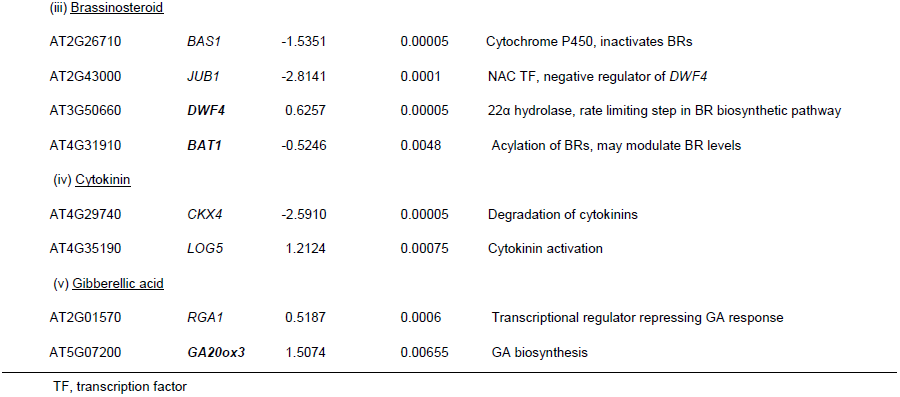
Selected genes differentially expressed in *Δppd*. Putative PPD2 targets are in bold.

| AGI Code | Gene Symbol | log<sub>2</sub> Fold Change | P value | Gene function |
| --- | --- | --- | --- | --- |
| *(A) Stomatal development* |  |  |  |  |
| AT1G05230 | *HDG2* | 0.5078 | 0.0016 | HD-ZIP IV protein, promotes stomatal development |
| AT1G12860 | *SCRM2* | 0.5127 | 0.0008 | bHLH TF, determinant of progression of stomatal cell lineage |
| AT1G26600 | *CLE9* | 0.8623 | 0.00225 | CLE peptide, possible role in meristemoid maintenance |
| AT1G34245 | *EPF2* | 1.1897 | 0.00005 | Secretory peptide, controls asymmetric cell division in stomatal lineage |
| AT1G80080 | ***TMM*** | 0.9809 | 0.00005 | Receptor, promotes asymmetric cell division in stomata development |
| AT3G26744 | *SCRM1* | 0.6699 | 0.00005 | bHLH TF, determinant of progression of stomatal cell lineage |
| AT3G57230 | *AGL16* | 0.4426 | 0.0055 | MADS box TF, promotes stomata from satellite meristemoids |
| AT5G07180 | *ERL2* | 1.1209 | 0.00005 | Receptor-like kinase, maintains stomatal stem cell activity |
| AT5G53210 | *SPCH* | 0.8734 | 0.00005 | bHLH TF, establishes stomatal cell lineage |
| AT5G60880 | *BASL* | 1.3827 | 0.00005 | Regulator of asymmetrically dividing stomatal-lineage cells |
| AT5G62230 | ***ERL1*** | 1.1739 | 0.00005 | Receptor-like kinase, maintains stomatal stem cell activity |
| *(B) Light signalling, circadian rhythm, and flowering genes.* |  |  |  |  |
| *(i) Light signalling* |  |  |  |  |
| AT2G24540 | ***AFR*** | -0.6003 | 0.0001 | F-box protein positive regulator of phyA signalling |
| AT2G42870 | *PAR1* | -0.6727 | 0.0028 | bHLH TF, negative regulator of shade avoidance |
| AT2G46340 | ***SPA1*** | 0.4398 | 0.0049 | Regulator of phyA signalling, repressor of photomorphogenesis |
| AT2G46970 | *PIL1* | -1.8117 | 0.00005 | Regulates light-mediated inhibition of hypocotyl elongation |
| *(ii) Circadian rhythm* |  |  |  |  |
| AT1G22770 | *GI* | 0.5658 | 0.0004 | Regulates circadian clock phyB signalling and flowering time |
| AT2G25930 | *ELF3* | 0.5306 | 0.0017 | Circadian clock component, interacts with phyB to control development |
| AT5G24470 | ***PRR5*** | 1.4340 | 0.00005 | Regulates circadian clock components and photomorphogenesis |
| AT5G59570 | ***BOA*** | 1.2499 | 0.00075 | Component of circadian clock |
| *(iii) Flowering* |  |  |  |  |
| AT5G47640 | ***NF-YB2*** | -0.7127 | 0.0019 | Interacts with CO, required for transcription of *FT* |
| AT5G60120 | ***TOE2*** | 0.4797 | 0.00165 | AP2 TF, represses *FT* expression |
| *(C) Hormone biosynthesis, catabolism, conjugation and signalling genes.* |  |  |  |  |
| *(i) Absciscic acid* |  |  |  |  |
| AT1G52400 | *BG1* | -2.1175 | 0.00005 | Regeneration of ABA from inactive ABA glucosides |
| AT4G04020 | ***FIB*** | -3.1345 | 0.00005 | Involved in ABA-mediated photoprotection |
| AT5G49450 | *bZIP1* | -1.1292 | 0.00005 | bZIP TF, activates ABA response genes |
| AT5G52310 | *RD29A* | -1.5379 | 0.00005 | Up-regulated by ABA |
| *(ii) Auxin* |  |  |  |  |
| AT1G70560 | *TAA1* | 1.2166 | 0.00005 | IAA biosynthesis |
| AT4G14550 | ***IAA14*** | -0.6555 | 0.0004 | Inhibits auxin regulated gene expression |
| AT4G27260 | ***GH3.5*** | -0.8659 | 0.00005 | IAA amido synthetase, alters free IAA levels |
| AT4G31500 | *SUR2* | -0.7410 | 0.00005 | Biosynthesis of glucosinolates, mutation confers auxin over production |
| (iii) Brassinosteroid |  |  |  |  |
| AT2G26710 | *BAS1* | -1.5351 | 0.00005 | Cytochrome P450, inactivates BRs |
| AT2G43000 | *JUB1* | -2.8141 | 0.0001 | NAC TF, negative regulator of *DWF4* |
| AT3G50660 | ***DWF4*** | 0.6257 | 0.00005 | 22α hydrolase, rate limiting step in BR biosynthetic pathway |
| AT4G31910 | ***BAT1*** | -0.5246 | 0.0048 | Acylation of BRs, may modulate BR levels |
| (iv) Cytokinin |  |  |  |  |
| AT4G29740 | *CKX4* | -2.5910 | 0.00005 | Degradation of cytokinins |
| AT4G35190 | *LOG5* | 1.2124 | 0.00075 | Cytokinin activation |
| (v) Gibberellic acid |  |  |  |  |
| AT2G01570 | *RGA1* | 0.5187 | 0.0006 | Transcriptional regulator repressing GA response |
| AT5G07200 | ***GA20ox3*** | 1.5074 | 0.00655 | GA biosynthesis |

TF, transcription factor

Other GO terms over represented in the dataset included those for “anthocyanin biosynthesis” and “light harvesting”. Notably, genes with key roles in anthocyanin biosynthesis; *CHS, CHI, FLS*, and *PAP1* were all down-regulated, as were light harvesting genes; *CAB2, LHCA6, LHCB2.3*, and *LHC4.3.* Since all of these genes are positively regulated by HY5^10^, putative HY5 binding sites^9^ were identified among genes in the *Δppd* dataset (Supplementary Table 1). HY5 was predicted to bind to 626 genes, a significant enrichment (22%) compared to the by chance prediction of 11.6%. About 21.6% of these HY5 targets in the dataset were up-regulated genes. There was also over representation of GO terms relating to pathogen defensive responses “defence response to fungus”, and “defence response to bacterium” in the dataset, with most of the genes down-regulated.

Both the *Δppd* seedling phenotype and the over-abundance of “light signalling response” GO terms in the RNA-Seq dataset suggested *PPD* genes might act as positive regulators of photomorphogenesis. To explore this possibility the dataset was scanned for genes involved in light signalling, circadian rhythm and the photoperiodic control of flowering time. Selected examples are given in Table IB. The up-regulation of *SPA1*, a negative regulator of phyA signalling and photomorphogenesis^12^, and the down-regulation of *AFR*, a positive regulator of phyA signalling^45^, both putative PPD2 targets, is consistent with PPD proteins having a direct and high level role in the modulation of light signalling. Furthermore, the long hypocotyl phenotype and down-regulation of *PAR1*^46^ and *PIL1*^47^ genes in *Δppd* would be expected if the mutant was defective in light signalling. These changes in the expression of light signalling genes in the *Δppd* mutant were confirmed by RT-qPCR analysis, with the opposite pattern of expression in *PPDOE* (Fig. 2b).

Regulators of the circadian clock that interact with phyB, such as *ELF3* and *GI*, and clock components *PRR5, BOA*, and *CHE*, were up-regulated (Table IB). However, these alterations in the transcription of circadian rhythm genes do not appear to explain the delayed flowering of *Δppd*. Expression of the key flowering time gene *FT*, is mainly up-regulated in long days by the transcriptional activator CO, and significantly, expression of some of the multiple regulatory components determining the activity of CO have been altered in *Δppd*. Examples include the up-regulation of *TOE2* and *SPA1*, and the down-regulation of *NF-YB2*, all putative targets of PPD2 (Table IB). Expression analysis confirmed the up-regulation of *ELF3, TOE2* and *BOA*, and down-regulation of *NF-YB2* in *Δppd* (Fig. 2b, Supplementary Fig. S9).

The presence of genes involved in hormone metabolism in the *Δppd* differential expression dataset suggested a possible role for PPD in the regulation of BR, auxin, cytokinin (CK), GA and ABA activities (Table IC). As an example of changes to gene expression known to increase BR levels, a key biosynthesis gene, *DWF4*^26^, was up-regulated, whereas genes involved in BR inactivation, *BAS1*^22^, *UGT73C5*^23^, and *BAT1*^32^, or reduced biosynthesis, *JUB1*^34^, were down-regulated. Similarly, the auxin biosynthesis gene *TAA1*, was up-regulated, whereas genes reducing auxin activity or response, *SUR2, IAA14, IAA3, GH3.3*, *GH3.5, GH3.17*, and genes for camalexin biosynthesis (*CYP71A12, CYP79B2, CYP79B3*), were all down-regulated. For CKs, a gene involved in activation, *LOG5*, was up-regulated, and a gene for degradation, *CKX4*, down-regulated. While the GA biosynthesis gene *GA20ox3*, and a GA responsive gene, *GASA14*, were both up-regulated, *RGA1*, a negative regulator of GA response was also up-regulated. In contrast to these examples a gene responsible for the regeneration of ABA from a pool of inactive ABA glucosides, *BG1*, and genes involved in ABA responses: *bZIP1, RD22, RD29A, KIN2, FIB, COR15a, COR413-PMI*, were all down-regulated. Since *bZIP1*, a key positive regulator of responses to ABA also activates genes involved in carbon and nitrogen metabolism^48^, the dataset was scanned for evidence of a reduction in expression of C- and N- metabolism genes. Interestingly, as would be expected for reduced bZIP1 activity, genes involved in C-metabolism (*BCA6, PPDK, ATBETAFRUCT4, BAM9, AS6, SWEET2, SWEET11, SWEET13, SWEET17, SUC2, SUC5*) and N-metabolism (*GDH2, TYROSINE AMINO TRANSFERASE, ASN1, IVD, PRODH1, BCE2, THA1, MCCA*) were all down-regulated in *Δppd*.

Differential expression in the mutant was confirmed for; *DWF4, BAS1, BAT1, JUB1, TAA1, SUR2, IAA3, IAA14, GH3.3, GH3.5, GA20ox3, GASA14, RGA1, BG1, bZIP1, FIB, RD29A, CKX4* and *LOG5* (Fig. 2c, Supplementary Fig. S9). Where examined, expression in the *PPDOE* genotype was generally the opposite of the deletion mutant or similar to wild type, the notable exception being for *GA20ox3*, where expression was up-regulated in both *Δppd* and *PPDOE*. These patterns of gene expression in the *Δppd* mutant are consistent with enhanced activities for BR, GA, auxin and CK, and reduced sensitivity to ABA.

Although it has been reported that auxins, CKs and GAs regulate vascular (pro)cambium cell proliferation^49^, the dataset was also scanned for putative direct targets of PPD that might explain the *Δppd* and *PPDOE* cambial cell proliferation phenotypes. A possible candidate, *ACL5*, was down-regulated. It has been reported that an *acl5* loss-of-function mutant has both increased vascular (pro)cambium cell proliferation and xylem cell differentiation^50^, characteristics similar to the *Δppd* mutant. Reduced expression of *ACL5* in the *Δppd* mutant was confirmed by RT-qPCR expression analysis (Fig. 2b).

**Multiple hormone responses are modulated by *PPD***. To confirm *PPD* genes modulate hormone activity, wild type, *Δppd*, and *PPDOE* genotypes were evaluated for responses to hormone and/or hormone biosynthesis inhibitor treatments (Fig. 3). Treatment with brassinazole (BRZ, an inhibitor of BR biosynthesis), reduced hypocotyl elongation in wild type seedlings grown in darkness, while treatment with the BR epibrassinolide (BL) reduced root growth in the light (Fig. 3a,b). The *Δppd* mutant was relatively insensitive to BRZ inhibition of hypocotyl elongation in the dark, but hyper-responsive to BL inhibition of root growth, while *PPDOE* had the opposite response. BRs also influence stomata development on the hypocotyl, with BL treatment increasing and BRZ decreasing the number of stomata (Fig. 3c). The *Δppd* mutant was hyper-responsive to BL stimulation and relatively insensitive to BRZ inhibition of stomata number, while in *PPDOE* there were reduced responses to both BL and the inhibitor. A similar set of experiments was used to determine responses to GA and the GA biosynthesis inhibitor paclobutrazol (PAC). Wild type seedlings responded to GA treatment with hypocotyl elongation and increased stomata number on the hypocotyl, whereas PAC treatment restricted both (Fig. 3d,e). Interestingly, hypocotyl elongation of both *Δppd* and *PPDOE* was hyper-responsive to GA and relatively insensitive to PAC, compared with wild type. Together with the up-regulation of a gene for GA biosynthesis in both genotypes, these responses suggest increased GA activity in *PPDOE* hypocotyls. However, the hyper-response to GA and insensitivity to PAC in stomata number on hypocotyls of *Δppd*, together with the relative insensitivity of *PPDOE* to either treatment, suggests PPD’s may also limit responses to GA (Fig. 3e). To assess auxin response wild type and *Δppd* seeds where germinated in the presence of 5 µM or 50 µM picloram (a synthetic auxin). Low concentrations of picloram stimulated and high concentrations inhibited hypocotyl elongation of the wild type. The increased sensitivity of *Δppd* to picloram observed is consistent with the mutant having an increase in endogenous auxin activity (Fig. 3f). Similarly, the hyper-sensitivity of *Δppd* to root growth inhibition by the synthetic cytokinin, 6-benzyl aminopurine (BAP), and the reduced response of *PPDOE*, suggests PPD’s also modulate endogenous cytokinin activity (Fig. 3g).

**Figure 3.**
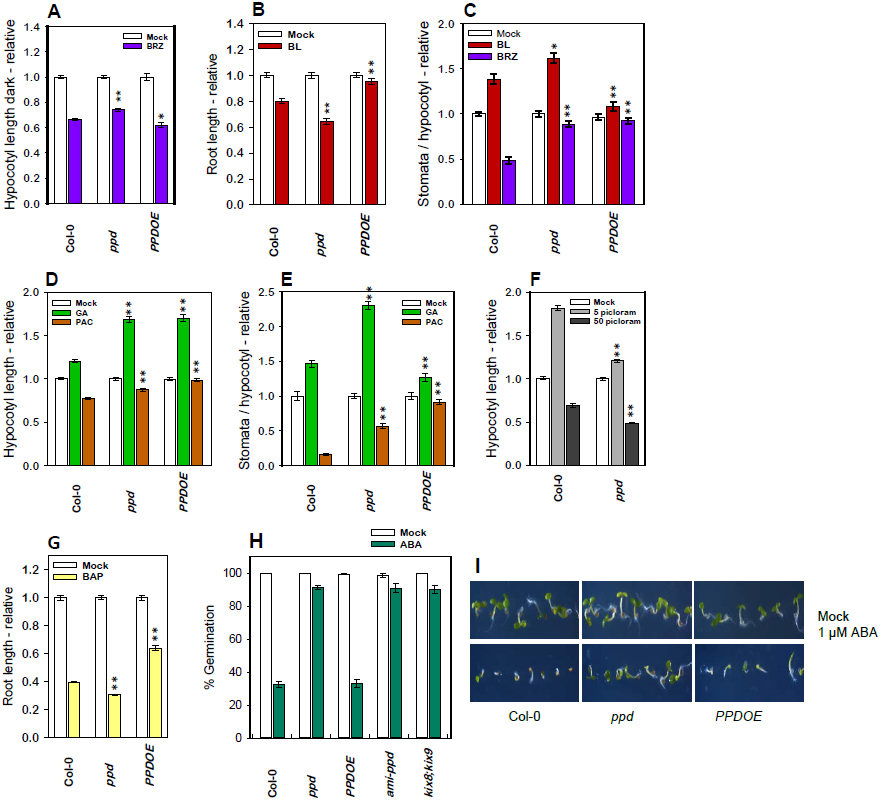
***PPD* modulates growth and developmental responses to hormones and inhibitors of hormone biosynthesis**. A-F Responses to hormone or hormone biosynthesis inhibitor treatments graphed as relative to mock treatments. (A) Dark grown hypocotyl length, 0.1µM brassinazole (BRZ) (n=20). (B) Root length, 1nM epibrassinolide (BL) (n=15). (C) Stomata per hypocotyl, 0.1µM BL, 1µM BRZ (n=30). (D) Hypocotyl length, 10µM gibberellic acid (GA), 1µM paclobutrazol (PAC) (n=35). (E) Stomata per hypocotyl, 10µM GA, 1µM PAC (n=30). Hypocotyl length, 5µM and 50µM picloram (n=35). (G) Root length, 1µM 6-benzylaminopurine (BAP) (n=18). Percentage seed germination at 3 days post stratification, 1µM abscisic acid (ABA). (I) Photographs taken at 4 dpg illustrating genotype differences in germination and greening in response to treatment with 1µM ABA. Significant differences from wild type \**P*<0.05 or \*\**P*<0.001(2-tailed Student’s *t*-test). Error bars represent ±SE.

As the gene expression profile in *Δppd* suggested a possible role for PPD’s in the modulation of ABA responses, genotypes with altered *PPD* complex expression were tested for sensitivity to ABA inhibition of seed germination (Fig. 3h,i). The insensitivity to ABA inhibition of seed germination of the *Δppd*, *ami-ppd* and *kix8;kix9* genotypes and the wild type like response of *PPDOE*, is consistent with PPD’s acting as positive regulators of ABA response.

***PPD* regulates BR biosynthesis**. Transgenic plants with overexpression of either the BR biosynthesis gene *DWF4* (*DWF4ox*)^51^ or the BR receptor *BRI1* (*BRI1ox*)^52^ have phenotypes similar to *Δppd*, with elongated hypocotyls, increased stomata numbers on hypocotyls, ABA insensitive germination and increased seed weights (Fig. 4a-d). In genetic interaction analyses *PPDOE* was epistatic to *DWF4ox* or *BRI1ox.* Furthermore, when *Δppd* was combined with a partial loss-of-function mutant of *DET2*^53^, a BR biosynthesis gene functioning upstream of *DWF4*, the double mutant (*det2;ppd*) had some increase in hypocotyl elongation, stomata numbers on hypocotyls, and insensitivity to ABA inhibition of seed germination, compared to *det2*. However, *det2* was epistatic to *Δppd* for both seed weight and the inhibition of hypocotyl elongation in dark grown seedlings (Fig. 4d-f). Collectively, these results are consistent with PPD’s acting as negative regulators of BR biosynthesis. RT-qPCR analysis indicated that there was no change in the expression of other genes involved in BR biosynthesis (*CPD, BR6ox2*), or signalling (*BIN2*) or transcriptional activation (*BES1, BZR1, BIM2, BEH2, BEH3*) in *Δppd* or *PPDOE* (Supplementary Fig. S9).

**Figure 4.**
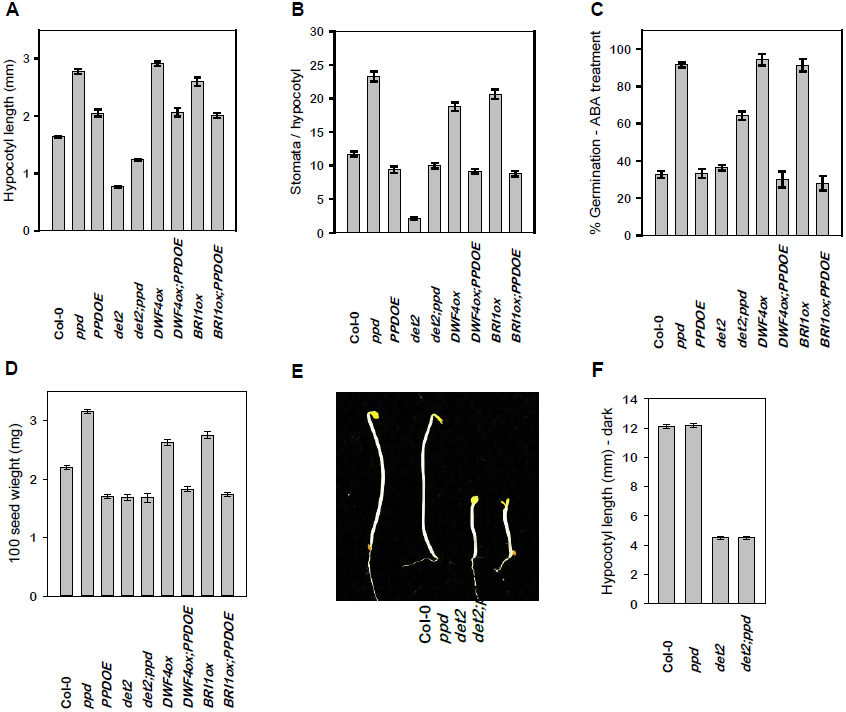
***PPD* is a negative regulator of BR biosynthesis.** Genetic analysis of interactions between genotypes with altered *PPD* function (*Δppd, PPDOE*) and genotypes with altered BR metabolism or signal transduction (*det2*, reduced biosynthesis, *DWF4ox*, increased biosynthesis, and *BRI1ox*, increased BR signalling). (A) Hypocotyl length, (n=30). (B) Stomata per hypocotyl, (n=40). (C) Percentage germination in the presence of 1µM ABA. (D) Seed weight. (E) Photograph of dark grown seedlings 4 dpg. (F) Hypocotyl length of dark grown seedlings (n=20). Error bars represent ±SE.

**Interactions between *PPD*, *CRY1* and *PHYB*, modulate light signalling.** Because transcriptomic analysis indicated PPD’s may regulate some of the initial steps in the light signalling cascade by repressing *SPA1* and activating *AFR* transcription, genetic interactions between *PPD* and *CRY1* or *PHYB* were evaluated. Seedlings of a *cry1* loss-of-function mutant had elevated stomata numbers on elongated hypocotyls and increased procambium cell proliferation in vascular tissues (Fig. 5b-d). Genetic interaction in the double *cry1;ppd* mutant was additive for hypocotyl length and stomata number on hypocotyls, but not for procambium cell proliferation where *cry1*, *Δppd* and the double mutant were all similar. For all these characteristics *PPDOE* was epistatic to *cry1*, suggesting that the reduced *SPA1* and/or elevated *AFR* expression in *PPDOE* (Fig. 2b) counteracted the defect in *CRY1* signalling. Seedlings of a *phyB* loss-of-function mutant also had increased stomata numbers on hypocotyls (Supplementary Fig. S7). Plants with overexpression of *phyB* (*phyBOE*), exhibited short hypocotyls, a reduction in stomata number on hypocotyls, early flowering, insensitivity to ABA inhibition of seed germination (Fig. 5), and reduced root growth (Supplementary Fig. S1). Procambium cell proliferation and seed weight of *phyBOE* was similar to wild type. Overexpression of *phyB* was epistatic to *Δppd* for most of the parameters examined; hypocotyl length, stomata numbers on hypocotyls, procambium cell proliferation, flowering time, seed weight, and root growth. However, leaf curvature, and the extended proliferation of meristemoids during leaf development where similar to *Δppd* (Fig 5a, Supplementary Fig.8). While the number of stomata on hypocotyls at 5 dpg was similar to *phyBOE* there was also extensive additional meristemoid cell proliferation on the hypocotyls of *phyBOE;ppd* seedlings. Hence, *phyB* overexpression counteracted most aspects of the *Δppd* mutant phenotype, but not the extended proliferation of stomatal lineage cells on hypocotyls, and leaves.

**Figure 5.**
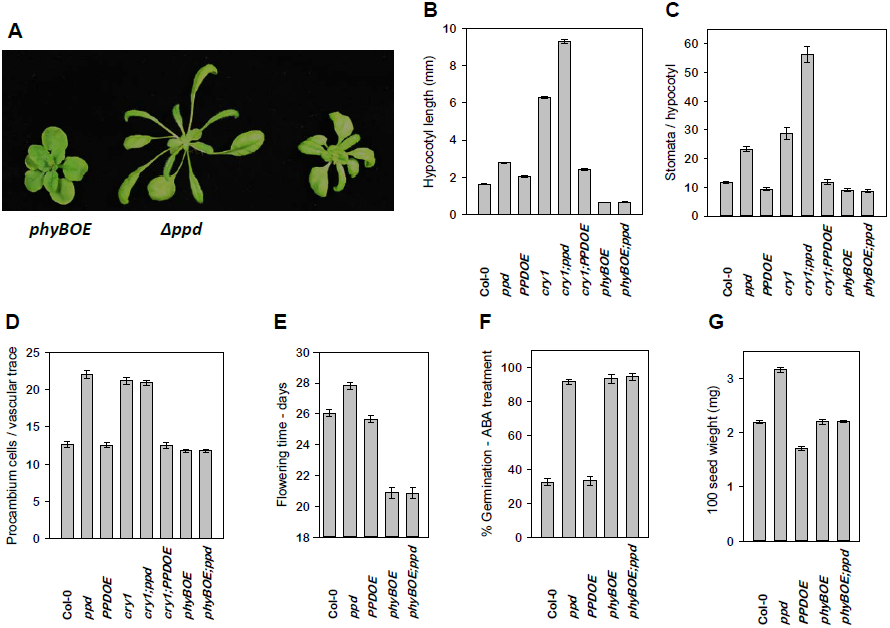
***PPD* genetically interacts with *CRY1* and *PHYB* to regulate light signalling.** (A) Comparison of *phyBOE* (over expression of phytochrome B), *Δppd*, and *Δppd;phyBOE* 21 dpg. (B) Hypocotyl length. (C)Stomata per hypocotyl. (D) Procambium cells per vascular trace section. (E) Flowering time. (F) Germination sensitivity to ABA. (G) Seed weight. Error bars represent ±SE.

The expression of a set of genes selected from the RNA-Seq *Δppd* dataset as indicators of; stomatal development, light signalling, floral induction, procambium cell proliferation and hormone metabolism, was compared between *Δppd, phyBOE*, and *phyBOE;ppd* (Fig. 6). For genes involved in light signalling (*SPA1, AFR, ELF3*) expression in the *phyBOE* and *phyBOE;ppd* genotypes was similar and opposite to the *Δppd* mutant. Expression of *TOE2* was down-regulated in *phyBOE*, up-regulated in *Δppd* and intermediate between the two in the double mutant, whereas expression of *NF-YB2* was down-regulated in all genotypes. Expression of *ACL5* was up-regulated in both *phyBOE* and *phyBOE;ppd*, in contrast to its down-regulation in the *Δppd* mutant. Unlike these examples, expression of *SPCH* was up-regulated in both *Δppd* and *phyBOE;ppd* but not in *phyBOE*, an expression pattern consistent with the meristemoid cell proliferation phenotypes (Supplementary Fig. S9).

**Figure 6.**
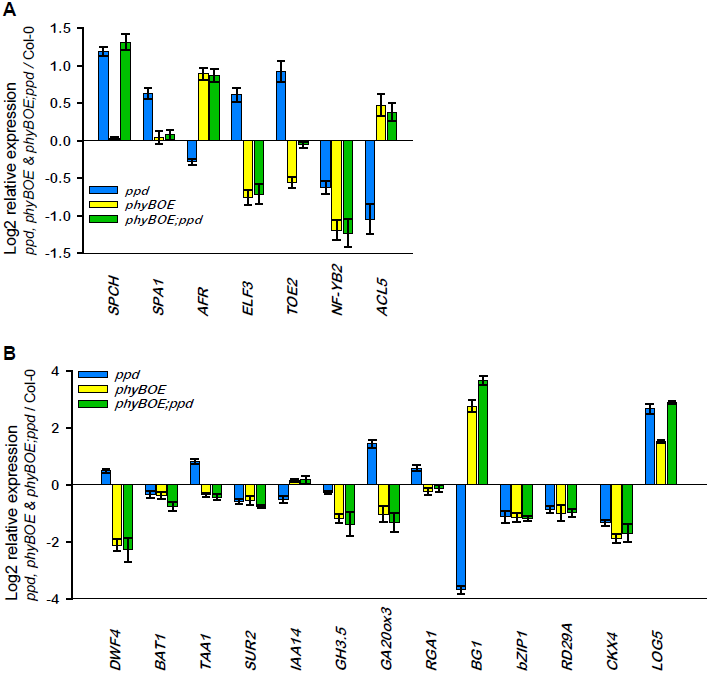
***PHYB* over expression alters the pattern of *Δppd* gene expression.** RT-qPCR expression analysis of selected genes regulating (A) stomatal development, light signalling, circadian rhythm, photoperiodic flowering, and cambium proliferation, and (B) hormone biosynthesis, catabolism, conjugation and signalling genes. Error bars represent ±SE.

Among the hormone biosynthesis genes examined (*DWF4, TAA1, GA20ox3*) expression in *phyBOE* and *phyBOE;ppd* was down-regulated, opposite to *Δppd* (Fig. 6b). Similarly, *phyB* overexpression altered transcription of the hormone signalling repressors, *RGA1* and *IAA14*. Expression of the ABA activation gene, *BG1* was also similar in the *phyBOE* and *phyBOE;ppd* genotypes and opposite to the deletion mutant. In contrast to this epistatic effect of *phyBOE* on the expression of genes involved in BR, auxin, GA and ABA biosynthesis or activation, the expression of genes controlling hormone inactivation (*BAT1, SUR2, GH3.5, CKX4*) was similar in all the genotypes. Hence, *phyBOE* probably counteracted the *ppd* loss-of-function influence on BR, auxin and GA activities by altering the biosynthesis of these hormones. Expression of the cytokinin activation gene *LOG5* was similar in *Δppd* and *phyBOE;ppd*, less in *phyBOE* seedlings but still up-regulated compared with wild type. Despite the possible increase in active ABA due to up-regulation of *BG1* in *phyBOE* and *phyBOE;ppd*, the expression of ABA response genes *bZIP1* and *RD29A* was similar in all genotypes. Collectively, results from genetic interactions between *CRY1/PHYB* and *PPD* and the pattern of gene expression in *phyBOE;ppd* plants, suggest the *PPD* genes act to modulate photomorphogenesis by regulating both the light signalling pathway and genes controlling hormone metabolism

**Discussion.** Increased leaf size resulting from reduced expression of the *PPD1* and *PPD2* genes in *Arabidopsis* has largely been attributed to the prolonged proliferation of meristemoids, precursor cells in the stomatal lineage. However, examination of *PPD* function in other species suggests there are alternative mechanisms for PPD control of organ size. In a recent report on the down-regulation of a *PPD* ortholog in legumes, enhanced general cell division and not altered meristemoid cell proliferation, appears to have caused increases in leaf and seed size^54^. Here I report a more extensive phenotype for the *Arabidopsis Δppd* mutant that reveals a role for the *PPD* genes as regulators of plasticity in growth and development throughout the life cycle. Features of the *Δppd* mutant phenotype include: tolerance to ABA inhibition of germination, hypocotyl elongation, increases in stomata number on hypocotyls, and (pro)cambium cell proliferation, delayed flowering time, and higher seed weight. Since the *kix8;kix9* mutant phenocopies the *Δppd* mutant, the PPD interacting KIX proteins are also likely to be a part of the molecular mechanism regulating these diverse aspects of plant development.

Whole-genome analysis of transcription identified PPD target genes regulating light signalling and hormone homeostasis that provide a possible explanation for many aspects of the complex mutant phenotype. The PPD complex may act to either repress or activate the transcription of genes, and in some cases both modes of action seem to be used to regulate antagonistic components of a particular signalling or biosynthesis pathway. Examples include the repression of *SPA1* and activation of *AFR* transcription to promote photomorphogenesis and the down-regulation of *DWF4* and up-regulation of *BAT1* to limit endogenous BR levels (Fig. 2b,c). Because light signalling influences many aspects of plant development by controlling endogenous phytohormone levels, it is possible that the *PPD* genes act only as positive regulators of the light signalling pathway. However, the *Δppd* mutant phenotype and its genetic interaction with *cry1* and *phyBOE* suggest there are some aspects of PPD action that are independent of light signalling. Although there are many similarities between the shade avoidance response and the *Δppd* phenotype; ABA insensitive seed germination, hypocotyl elongation, petiole elongation, down-regulation of anthocyanin biosynthesis and pathogen defence genes, and increased activity of the phytohormones, auxin, BR, GA, CK, other aspects of the mutant phenotype; delayed flowering, increased leaf and seed size and higher leaf stomatal density, are opposite to those expected from plants growing in shade.

The *PPD* genes may also use a combination of direct and indirect (via light signalling) mechanisms to regulate specific developmental pathways. One example is the increased (pro)cambium proliferation of the *Δppd* mutant. Although the pathways controlling cambial cell proliferation are not fully understood it has been shown that the auxin dependent bHLH transcription factor TMO5 forms a heterodimer complex with another bHLH protein, LHW, and this TMO5-LHW complex directly up-regulates transcription of the CK biosynthesis genes *LOG3 and LOG4*, resulting in a CK mediated increase in (pro)cambium cell proliferation^55^. Thermospermine produced by ACL5 represses the translational inhibitory effect of upstream ORFs located in the 5ʹ leader sequence of *SACL* genes. These genes encode bHLH transcription factors that compete with TMO5 for its LHW binding partner, thereby reducing the amount of active TMO5-LHW^56–57^. The *PPD* genes act as positive regulators of *ACL5* transcription (Fig. 2b), and thereby may limit formation of the TMO5-LHW complex and cambial cell proliferation. Interestingly, light signalling also influences vascular growth. Loss-of-function mutants for *phyB* and *hy5* have fewer xylem vessels in stem vascular than wild type^58–59^. This influence of the phytochromes on vascular growth may contribute to increased fitness in an exposed environment, since cucumber seedlings deficient in phyB grown outdoors exhibited low survival due to hypocotyl fracture^58^. Surprisingly then, defects in blue light, cryptochrome-mediated, signalling resulted in increased procambium cell proliferation in seedlings (Fig. 5). Since reduced cryptochrome signalling occurs in plants grown in shade, increased amounts of vascular tissue may be produced under these conditions to provide mechanical strength for elongating hypocotyls and petioles. The blocking of increased procambium cell proliferation in *cry1* by *PPDOE* and in *Δppd* by *phyBOE* (Fig. 5d) suggest that a mutually antagonistic interaction between *PPD* and light signalling modulates the rate of cambial cell proliferation. Because auxin activates *TMO5* expression, and light signalling regulates the level of endogenous auxin, the effect of *PPD* expression on cambial cell proliferation may result from a combination of direct control of *ACL5* transcription and indirect regulation of auxin homeostasis via modulation of light signalling.

A mutually antagonistic interaction between *PPD* and light signalling also regulates stomatal development on the hypocotyl (Fig. 5c). While it is known that stomata number on *Arabidopsis* hypocotyls is increased by BR or GA^60–62^ (Fig. 3), the influence of cryptochrome and phytochrome-mediated light signalling on hypocotyl stomatal development is unknown. Here I report that both cry1 and phyB act to limit stomata number on the hypocotyl. Overexpression of *PPD1* blocked the increased stomata of a *cry1* loss-of-function mutation, while *phyB* over-expression blocked the increase in stomata number on the hypocotyl of the *Δppd* mutant (Fig. 5c). The stomatal phenotype of *phyBOE;ppd* hypocotyls, which have extensive meristeomoid proliferation but no increase in stomata number, suggests excess activated phyB acts to inhibit stomatal development on hypocotyls downstream of PPD. Since over expression of *phyB* likely limits BR and GA hormone levels and *Δppd* deletion results in increased *SPCH* expression and a higher population of meristemoid cells in the epidermis, BR and GA maybe required for the development of the proliferating meristemoids into stomata. Collectively, these results suggest the intriguing possibility that seedlings respond to shading with a coordinate increase in hypocotyl length, stomata production and vascular growth, and that PPD provides a braking mechanism to prevent excessive developmental response to shading.

There are numerous possible explanations for the delayed flowering phenotype of the *Δppd* mutant. Expression of the flowering time gene *FT* is mainly up-regulated in long days by the transcriptional activator CO, and significantly, expression of some of the multiple regulatory components determining the activity of CO have been altered in the *Δppd* mutant. Examples include the up-regulation of *TOE2* and the down-regulation of *NF-YB2*, both putative targets of PPD2 (Table II). TOE2 forms a complex with CO to suppress CO activation of *FT*^63^, while NF-YB2 is a part of the NF-Y complex that enhances the binding of CO to the *FT* promoter^64^. Other changes in the *Δppd* mutant that could lead to reduced CO activity were the increased expression of *SPA1*, since COP1-SPA1 complexes degrade CO^65^, and up-regulation of *RGA1*^66^. It is possible that a combination of these factors determine the delayed flowering of the *Δppd* mutant since expression of *TOE2, SPA1* and *RGA1* were reduced in *PPDOE* (Fig. 2). The down-regulation of *NF-YB2* in *Δppd*, *phyBOE* and *phyB;ppd*, together with *phyBOE* epistasis (Fig. 6), suggests that reduced expression of *NF-YB2* did not significantly contribute to delayed flowering of the *Δppd* mutant.

Since both *ppd* and *kix8;kix9* loss-of-function genotypes are insensitive to the inhibitory effect of ABA on seed germination (Fig. 3) the PPD complex maybe required for activation of the ABA signalling pathway. Light signalling positively regulates ABA response by HY5 binding to the promoter of *ABI5*, a bZIP transcription factor controlling the expression of genes responsible for the inhibition of seed germination^67^. Although PPD might influence ABA responses indirectly by modulating light signalling^68^, the results of gene expression profiling suggest an alternative molecular mechanism involving regulation of *bZIP1*, which encodes a transcription factor controlling many ABA response genes. Both *bZIP1*, and *RD29A* which it activates, were down-regulated in *Δppd, phyBOE*, and *phyBOE;ppd* (Fig. 6), all genotypes tolerant to the inhibitory effect of ABA on seed germination (Fig. 5). This expression pattern may reflect a cascade effect, since *bZIP1* is activated by the NF-Y complex, and NF-Y complex function is likely to be reduced in *Δppd, phyBOE* and *phyBOE;ppd* due to the down-regulation of *NF-Y2*.

It has been reported that in *Arabidopsis* brassinosteroids modulate the transcription of genes for seed development specific pathways and are required for the development of normal seed size and weight^69^. Furthermore, overexpression of *DWF4* results in increased seed weight^51^. Here I report increases in seed weight for *Δppd, kix8;kix9*, and *BRI1ox* genotypes. In genetic interaction analysis *PPDOE* was epistatic to *DWF4ox* and *BRI1ox*, whereas *phyBOE* was epistatic to *Δppd* (Figs. 4d & 5g). These results confirm a role for BR in the regulation of seed weight, and suggest that PPD limits growth of developing seeds by restricting BR biosynthesis, either directly or by modulating light signalling. Collectively, these results reveal a previously unrecognised role whereby PPD acts as a braking mechanism to limit developmental plasticity. Because *PPD* modulates developmental traits of economic importance there is potential to use tissue-specific alterations in *PPD* expression to obtain improvements in yield and stress tolerance in plants.

## Methods

**Plant Material.** The *Arabidopsis Δppd* mutant and transgenic *Δppd::PPD1* complemented, and *Δppd::PPD1* over-expression (*PPDOE*) genotypes in a Landsberg *erecta* (L*er*) ecotype background have been described previously^38^. All other *Arabidopsis* plants used in this study are in the Columbia (*Col-0*) background. The *Δppd* mutant and transgenic *Δppd::PPDOE* genotypes were backcrossed with wild-type *Col-0* for four generations to obtain *Δppd Col-0* and *PPDOE Col-0*. Mutants *det2-1* (CS6159), *cry1* (CS6955), and a transgenic phyB over expression genotype, *phyBOE* (35S::*phyB*, CS3081) were obtained from the Arabidopsis Biological Resource Center. Transgenic lines with over expression of *DWF4* (35S::*DWF4ox)* or *BRI1* (35S::*BRI1ox*) as previously described^52^, were supplied by Joanne Chory. The transgenic *PPD* silencing line *ami-ppd* and the *kix8;kix9* double mutant^39^, were supplied by Dirk Inzé. The *det2, DWF4ox, BRI1ox, cry1*, and *phyBOE* genotypes were crossed with *Δppd* or *PPDOE* and double mutants (*det2;ppd, cry1;ppd, phyBOE;ppd*) or triple mutants (*DWF4ox;ppd::PPDOE, BRI1ox;ppd::PPDOE*, and *cry1;ppd::PPDOE*) selected from F_2_ populations. A combination of phenotype and genotyping by PCR with gene specific primers (Supplementary Table S2) identified homozygous lines.

**Measurement of Flowering Time.** Seeds were stratified at 4 ^0^C for 3 days in the dark then germinated and plants grown in a soil mix in a controlled environment growth room at 22 ^0^C, 65% relative humidity, with 14 h white light/10 h dark cycles. Light intensity was 200 μm·m^-2^·s^-1^ from a combination of Sylvania GRO-LUX F36W/GRO-T8 and Phillips TLD 58W/840 fluorescent tubes. Flowering time was recorded as the number of days post germination taken for inflorescences to attained 1 cm in height. The experiment was replicated twice with 10 plants per genotype in each experiment.

**Seed Weight Measurements.** The weight of batches of 100 seeds of each genotype, were determined using a Mettler Toledo analytical balance. Results are the averages of seven replicates.

**Seedling Growth Conditions.** For hypocotyl growth, stomatal density, cotyledon vascular procambium cell proliferation, hormone response, RNA Seq and quantitative RT-PCR gene expression experiments, seed were surface sterilised with 70% ethanol, 0.01% Triton X-100 for 10 min, followed by 100% ethanol for 5 min, air dried on sterile filter paper, and transferred to media plates containing half-strength MS salts and vitamins (Duchefa Biochemie), 1% sucrose and 0.8% agar. Plates were then incubated for 3 days at 4 ^0^C in the dark and grown at 24 ^0^C under a 14 h light/10 h dark daily cycle. White light was provided by fluorescent tubes (Philips TLD 58W/840) at an intensity of 100 μm·m^-2^·s^-1^.

**Microscopy.** Hypocotyl length and hypocotyl stomatal density were scored on plate grown seedlings sampled 5 days post germination (dpg), cleared in 85% lactic acid, for 1 min at 101 ^0^C, 15 psi, and mounted in the same solution on microscope slides. Total number of stomata on each hypocotyl was counted with an Olympus BX50 microscope using interference contrast (n=40), while hypocotyl length was measured using a Leica Wild M3Z binocular microscope fitted with an eyepiece micrometer (n=30). For observations on cotyledon stomatal development seedlings were sampled at 10 dpg and cleared by passaging through the following solutions: 70% ETOH, 70% ETOH/10% acetic acid, 1 ETOH:1 2.5M NaOH, each for 1 hr, then overnight in 85% lactic acid. Whole plant images were obtained using an Olympus SZX12 binocular microscope and DP20 digital camera. Dark grown hypocotyl length was measured on seedlings sampled 4 dpg after incubation in continuous darkness at 24^0^C. For electron microscopy, hypocotyls sampled at 3 dpg were observed directly using an FEI Quanta 200 environmental scanning electron microscope.

**Histology.** Plate grown seedlings sampled 10 dpg were fixed in 4% glutaraldehyde, 1% formaldehyde, in 50 mM phosphate buffer, pH 7, overnight, embedded in Technovit 7100 resin (Heraeus Kulzer) following the manufacturers protocol, and 7 µm sections prepared using a Lieca RM2045 microtome. Sections were stained with Toluidine blue O, the number of procambium cells per cotyledon petiole vascular trace transverse section recorded (n=30) and digital images obtained with an Olympus BX50 microscope analySIS^B^ system.

**Transcript Analysis.** Total RNA was extracted from ten-day-old seedlings grown on 0.5 MS medium at mid-morning in the light cycle using an RNeasy Plant Kit (Qiagen) and genomic DNA contamination eliminated by on-column DNase I digestion using an RNase-Free DNase Set (Qiagen). RNA-Seq library preparation, high-throughput sequencing and data analysis services were provided by Oxford Gene Technology. RNA quality of the samples was determined using a 2100 Bioanalyzer (Agilent), and mRNA libraries prepared with the Illumina TruSeq RNA Sample Prep Kit v2. Sample sequencing was performed on an Illumina HiSeq2000 platform using TruSeq v3 chemistry. An average of 55,073,058 paired end reads was obtained per sample. Reads were mapped to the TAIR10 *Arabidopsis* genome assembly using Bowtie version 2.02, splice junctions identified using TopHat version 2.09 and Cufflinks version 2.1.1 was used to perform transcript assembly, abundance estimations and differential expression between wild-type and *Δppd* samples.

For real time reverse transcription quantitative PCR analysis first-strand complementary DNA was synthesised from 1µg total RNA, purified as described above, using both random hexamer and oligo(dT)_18_ primers and Maxima Reverse Transcriptase (Thermo Scientific) according to the manufacturer’s protocol. Quantitative PCR analyses were undertaken using a LightCycler 480 real-time PCR instrument (Roche), LightCycler 480 SYBR Green Master reagents version 12 (Roche), and gene-specific oligonucleotide primers (Supplementary Table S3). Transcript abundance was calculated using LightCycler 480 instrument software based on the ΔΔCT-method and relative expression of a gene was calculated from the ratio of test genotype samples to the reference Col-0 wild-type. Both *actin2* and *tubulin* were used to normalise different samples. Four biological replicates and at least four technical replicates were used for each genotype.

**Seedling Hormone-response Growth Assays.** Epibrassinolide (Santa Cruz Biotechnology), and brassinazole (Santa Cruz Biotechnology) dissolved in dimethlsulphoxide (DMSO), gibberellic acid (ACROS organics) and picloram (Sigma) in ethanol, paclobutrazol (Sigma) in acetone, abscisic acid in methanol and 6-benzylaminopurine (Sigma) in 0.1 N KOH, were filter sterilised and incorporated into 0.5 MS media plates. DMSO, ethanol, methanol or acetone as appropriate, were used for mock treatments. Seedlings were grown in light for 5 days before hypocotyl stomatal density (n=30), hypocotyl length (n=35) or root length (n=15) at 4 days or percentage germination (n=200) at 3 days post germination were analysed. Hypocotyl length (n=20) of dark grown seedlings was analysed at 4 days post germination. Each treatment was repeated twice.

## Acknowledgements

This work was funded by grants from Royal Society of New Zealand Marsden Fund (AGR304), Foundation for Research, Science & Technology (C10X0816) and Cambium Genetics. I thank the Manawatu Microscopy & Imaging Centre for the use of microscopy resources.

## Competing financial interests

The author declares no competing financial interests.

